# Silicon Nitride Inactivates SARS-CoV-2 *in vitro*

**DOI:** 10.1101/2020.08.29.271015

**Authors:** Caitlin W. Lehman, Rafaela Flur, Kylene Kehn-Hall, Bryan J. McEntire, B. Sonny Bal, Ryan M. Bock

**Author notes:** Corresponding Author: Ryan M. Bock.

## Abstract

**Introduction:** Severe acute respiratory syndrome coronavirus 2 (SARS-CoV-2), which is responsible for the COVID-19 pandemic, remains viable and therefore potentially infectious on several materials. One strategy to discourage the fomite-mediated spread of COVID-19 is the development of materials whose surface chemistry can spontaneously inactivate SARS-CoV-2. Silicon nitride (Si_3_N_4_), a material used in spine fusion surgery, is one such candidate because it has been shown to inactivate several bacterial species and viral strains. This study hypothesized that contact with Si_3_N_4_ would inactivate SARS-CoV-2, while mammalian cells would remain unaffected.

**Materials:** SARS-CoV-2 virions (2×10^4^ PFU/mL diluted in growth media) were exposed to 5, 10, 15, and 20% (w/v) of an aqueous suspension of sintered Si_3_N_4_ particles for durations of 1, 5, and 10 minutes, respectively. Before exposure to the virus, cytotoxicity testing of Si_3_N_4_ alone was assessed in Vero cells at 24 and 48 hour post-exposure times. Following each exposure to Si_3_N_4_, the remaining infectious virus was quantitated by plaque assay.

**Results:** Vero cell viability increased at 5% and 10% (w/v) concentrations of Si_3_N_4_ at exposure times up to 10 minutes, and there was only minimal impact on cell health and viability up to 20% (w/v). However, the SARS-CoV-2 titers were markedly reduced when exposed to all concentrations of Si_3_N_4_; the reduction in viral titers was between 85% - 99.6%, depending on the dose and duration of exposure.

**Conclusions:** Si_3_N_4_ was non-toxic to the Vero cells while showing strong antiviral activity against SARS-CoV-2. The viricidal effect increased with increasing concentrations of Si_3_N_4_ and longer duration of exposure. Surface treatment strategies based on Si_3_N_4_ may offer novel methods to discourage SARS-CoV-2 persistence and infectivity on surfaces and discourage the spread of COVID-19.

## 1. INTRODUCTION

The spread of viruses through contaminated surfaces is of concern in crowded indoor environments such as schools, nursing homes, hospitals, and day-care centers.^1^ In addition to the aerosol transmission of respiratory viruses due to sneezing, coughing, and talking, evidence suggests that contaminated hands and fomites also play a significant role in the person-to-person proliferation of disease.^2^ The global spread of COVID-19, the disease caused by the coronavirus designated as SARS-CoV-2, has rekindled interest in the role of contaminated surfaces in viral transmission.

Viruses can survive for long periods on a variety of surfaces. Rotavirus is the agent that causes acute diarrhea in pediatric patients. In fecal samples, rotavirus particles are stable even after 2½ months of storage at temperatures above 30°C, with even longer survival times at lower temperatures.^3^ The 2002 outbreak of severe acute respiratory syndrome (SARS) in China was caused by SARS-CoV;^4, 5^ a virus that is infective for up to 9 days in suspension, and 6 days in the dried state.^6^ The pathogenic coronavirus HuCoV-229E causes upper respiratory infections in humans; at room temperature, this virus can persist for up to 5 days on surfaces ranging from glass to stainless steel.^7^ A recent report has shown survival times of 4 to 72 hours of SARS-CoV-2 on plastic, copper, and cardboard surfaces, and up to 7 days on surgical masks.^8^

Surface decontamination by disinfectant application,^9^ or by irradiation,^10, 11^ can control fomite-induced viral spread. Some materials, notably metals such as copper (Cu), zinc (Zn), iron (Fe), and silver (Ag) have also been shown to rapidly inactivate viruses, including SARS-CoV-2.^7, 12, 13^ Incorporation of Cu alloy surfaces in health care facilities reduces the microbial burden and the incidence of nosocomial infections.^14–16^ Quaternary ammonium compounds such as ammonium chloride are also capable of inactivating viruses by deactivating the protective lipid coating that envelopes viruses like SARS-CoV-2 rely on; these compounds are used as disinfectants to clean surfaces and remove persistent viral particles.^17^

Silicon nitride (Si_3_N_4_) is a non-oxide ceramic that is FDA-cleared for use in implantable spinal fusion devices; clinical data have shown excellent long-term outcomes in both lumbar and cervical fusion with Si_3_N_4_ when compared to other spine biomaterials such as bone grafts, titanium, and polyetheretherketone.^18–23^ In aqueous environments, Si_3_N_4_ spine implants undergo surface hydrolysis, resulting in the microscopic elution of ammonia which is converted into ammonium, nitrous oxide, and other reactive nitrogen species that inhibit bacterial growth and proliferation.^24^ This inherent resistance to bacterial colonization may explain the lower incidence of bacterial infection with Si_3_N_4_ spinal implants (< 0.006%) when compared to other spine biomaterials (2.7% to 18%).^25^

Similar to bacterial inhibition, viral exposure to an aqueous solution of Si_3_N_4_ particles inactivated H1N1 (Influenza A/Puerto Rico/8/1934), Enterovirus (EV-A71), and Feline calicivirus.^26^ Recent findings have shown that Si_3_N_4_ particles in aqueous suspensions also inactivated SARS-CoV-2 with antiviral activity that compared favorably to Cu ions. Unlike Si_3_N_4_, Cu exhibited toxicity to mammalian cells.^12^ The present study hypothesized that exposure to Si_3_N_4_ would not elicit a toxic response from mammalian cells under experimental conditions while demonstrating a time- and dose-dependent inactivation of SARS-CoV-2.

## 2. MATERIALS AND METHODS

### 2.1. Preparation of Si_3_N_4_ powder

A doped Si_3_N_4_ powder (SINTX Technologies, Inc., Salt Lake City, UT USA) with a nominal composition of 90 wt.% *α*-Si_3_N_4_, 6 wt.% yttria (Y_2_O_3_), and 4 wt.% alumina (Al_2_O_3_) was prepared by aqueous mixing and spray-drying of the inorganic constituents, followed by sintering of the spray-dried granules (~1700°C for ~3 h), hot-isostatic pressing (~1600°C, 2 h, 140 MPa in N_2_), aqueous-based comminution, and freeze-drying.^27^ The resulting powder had a trimodal distribution with an average particle size of 0.8 ± 1.0 μm as shown in Figure 1. Doping Si_3_N_4_ with Y_2_O_3_ and Al_2_O_3_ is essential to densify the ceramic and convert it from its *α*- to *β*-phase during sintering. The mechanism of densification is via dissolution of *α*-phase and subsequent precipitation of *β*-phase grains facilitated by the formation of a transient intergranular liquid that solidifies during cooling. *β*-Si_3_N_4_ is therefore a composite composed of about 10 wt.% intergranular glass phase (IGP) and 90 wt.% crystalline *β*-Si_3_N_4_ grains.

**Figure 1.**
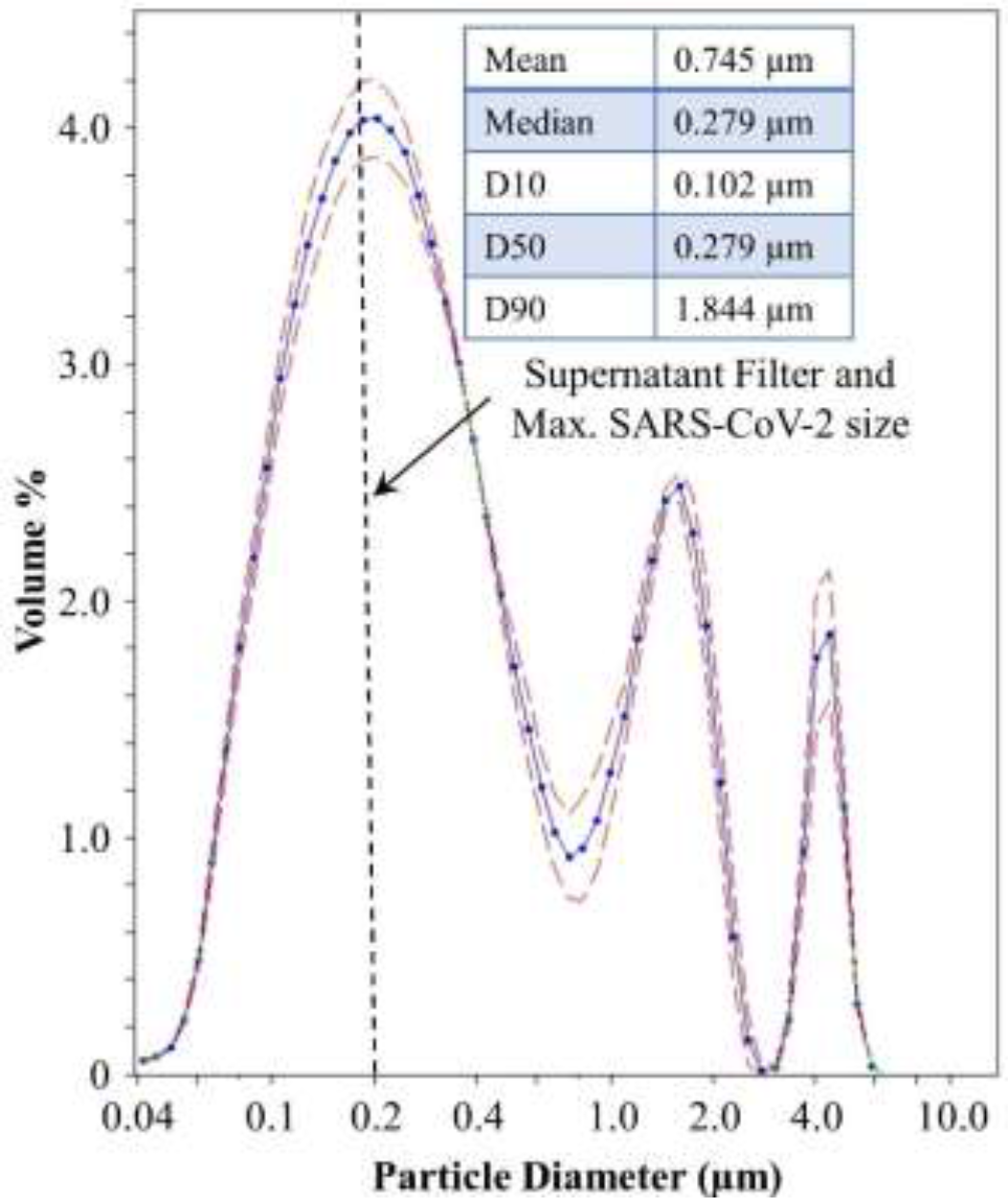
Silicon nitride powder particle size distribution.

### 2.2. Cell culture and viral stocks

Vero green African monkey kidney epithelial cells (ATCC, Cat #CCL-81) were chosen for this analysis due to their ability to support high levels of SARS-CoV-2 replication and their use in antiviral testing.^28, 29^ These cells were cultured in DMEM (VWR, Cat # 10128-206) supplemented with 10% FBS (VWR Cat # 97068-085), 1% L-glutamine (VWR Cat # 45000-676), and 1% penicillin/streptomycin (VWR Cat # 45000-652). Cells were maintained at 37°C and 5% CO_2_. SARS-CoV-2 isolate USA-WA1/2020 was obtained from BEI Resources (Cat #NR-52281). Vero cells were inoculated with SARS-CoV-2 (MOI 0.1) to generate viral stocks. Cell-free supernatants were collected at 72 hours post-infection and clarified via centrifugation at 10,000 rpm for 10 minutes and filtered through a 0.2 μm filter (Pall, Cat # 4433). Stock virus was titered according to the plaque assay protocol detailed below.

### 2.3. Cell viability assays

The Si_3_N_4_ powder was suspended in 1 mL DMEM growth media in microcentrifuge tubes. Tubes were vortexed for 30 seconds to ensure adequate contact and then placed on a tube revolver for either 1, 5, or 10 minutes. At each time point, the samples were centrifuged, and the supernatant was collected and filtered through a 0.2 μm filter. Clarified supernatants were added to cells for either 24 or 48 hours. Untreated cells were maintained alongside as controls. Cells were tested at each time point using CellTiter Glo (Promega, Cat #G7570), which measures ATP production, to determine cell viability.

### 2.4. Antiviral activity testing

SARS-CoV-2 was diluted in DMEM growth media to a concentration of 2 x 10^4^ PFU/mL. Four mL of the diluted virus was added to tubes containing silicon nitride at 20, 15, 10, and 5% (w/v). The virus without Si_3_N_4_ was processed in parallel as a control. Tubes were vortexed for 30 seconds to ensure adequate contact and then placed on a tube revolver for either 1, 5, or 10 minutes, while a virus only control was incubated for the maximum 10 minutes. At each time point, the samples were centrifuged, and the supernatant was collected and filtered through a 0.2 μm filter. The remaining infectious virus in the clarified supernatant was quantitated by plaque assay. An overview of the antiviral testing method is provided in Figure 2.

**Figure 2.**
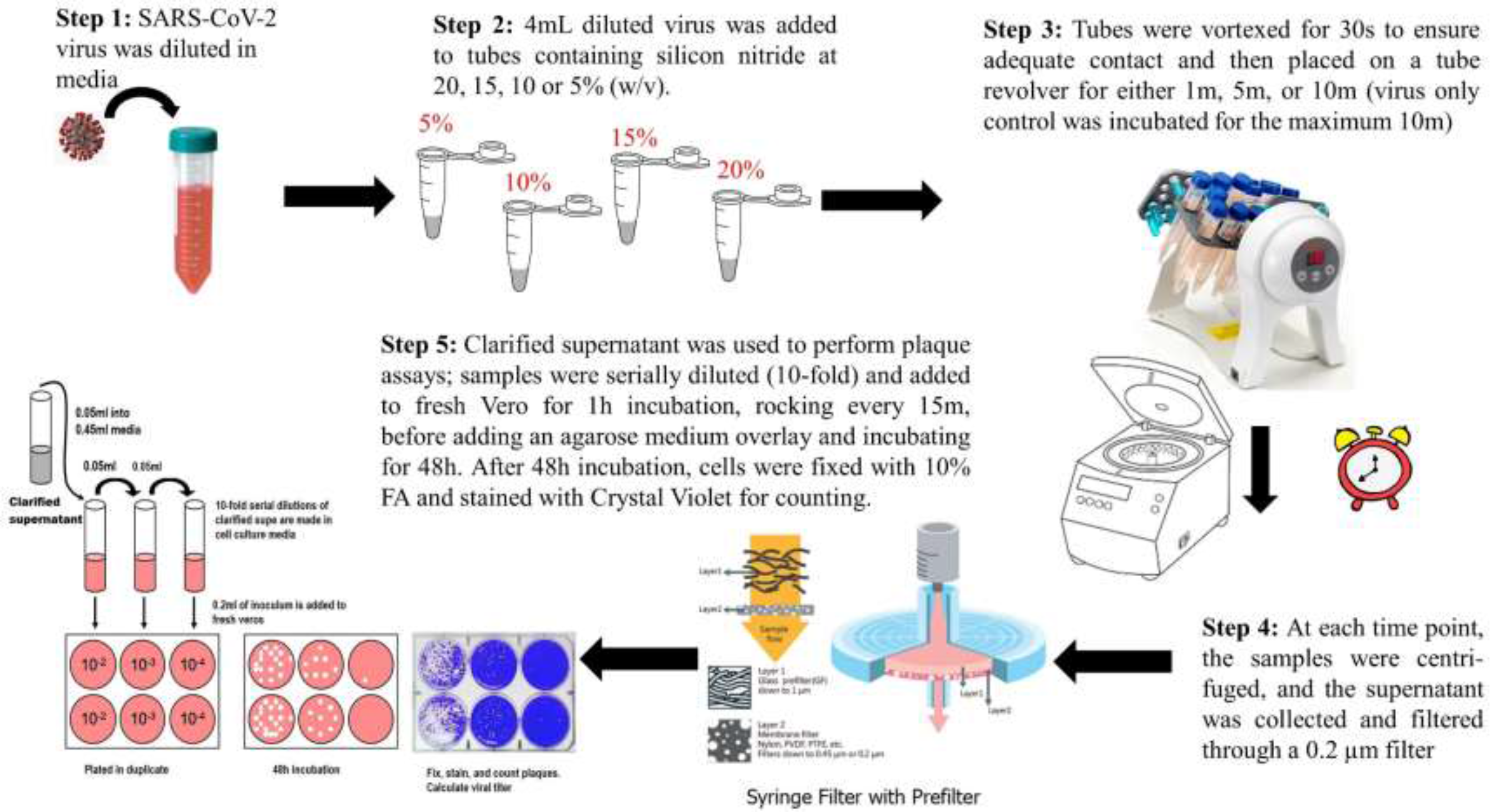
A Pictorial overview of the antiviral testing method.

### 2.5. SARS-CoV-2 plaque assay

Vero cells were plated at 2 x 10^5^ cells/well in a 12-well plate on the day before the plaque assay. Clarified supernatants from the antiviral testing were serially diluted (10-fold) and 200 μL was added to Vero cells which were incubated for 1 hour at 37°C, 5% CO_2_. Plates were rocked every 15 minutes to ensure adequate coverage and at 1 hour, a 1:1 ratio of 0.6% agarose and 2X EMEM (VWR, Cat # 10128-758) supplemented with 5% FBS, 2% penicillin/streptomycin, 1% non-essential amino acids (VWR, Cat # 45000-700), 1% sodium pyruvate (VWR, Cat # 45000-710), and 1% L-glutamine was added to the cells before incubating for 48 hours at 37°C, 5% CO_2_. After incubation, the cells were fixed with 10% formaldehyde and stained with 2% crystal violet in 20% ethanol for counting.

## 3. RESULTS

### 3.1. No toxicity of Si_3_N_4_ to Vero cells up to 20 wt.%/vol

The impact of Si_3_N_4_ on eukaryotic cell viability was tested. Si_3_N_4_ was resuspended in cell culture media at 5, 10, 15, and 20% (w/v). Samples were collected at 1, 5, and 10 minutes and added to Vero cells. Vero cell viability was measured at 24 and 48 hours post-exposure (Figure 3A and 3B). No significant decrease in cell viability was observed at either 24 or 48 hours post-exposure with 5%, 10%, or 15% silicon nitride. A small impact on cell viability (~10% decrease) was observed at 48 hours in cells exposed to 20% Si_3_N_4_. Interestingly, a ~10% increase in Vero cell viability was observed at 48 hours with the 5% - 10 minute and 10% - 10 minute samples (Figure 3B), suggesting that Si_3_N_4_ may be stimulating cell growth or cellular metabolism under these conditions. These data indicate that Si_3_N_4_ has minimal impact on Vero cell health and viability up to 20 wt.%/vol.

**Figure 3.**
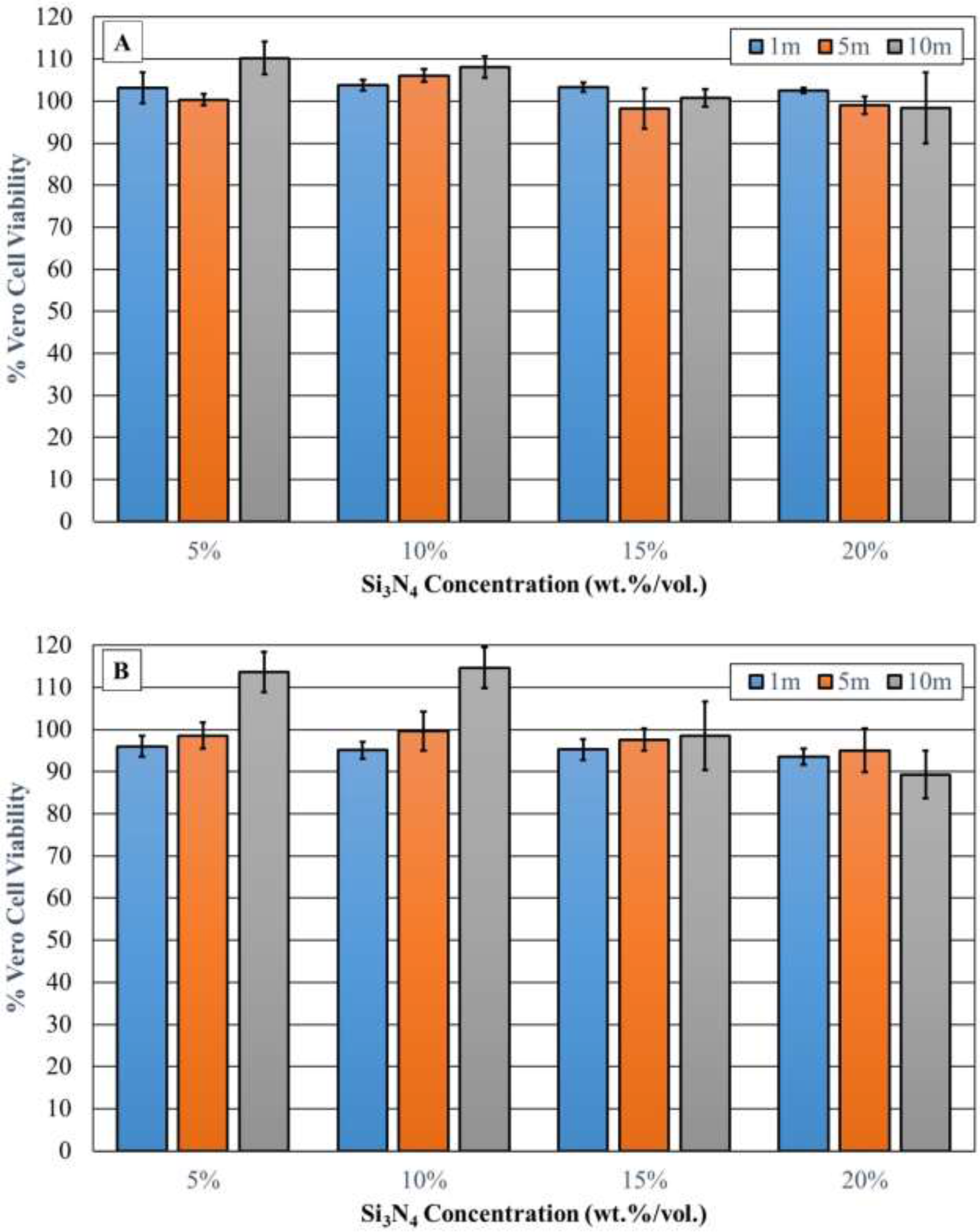
Cytotoxicity testing of silicon nitride. Silicon nitride at concentrations of either 5, 10, 15 or 20 wt.%/vol (n=4) were incubated with cell culture media for 1, 5, and 10m. At each timepoint, the samples were centrifuged, and the supernatant was collected and filtered through a 0.2um filter. The clarified supernatant was added to Vero cells and their viability assessed at 24 h (A) and 48 h (B). Untreated Vero cells served as a control and were set to 100% viability.

### 3.2. Si_3_N_4_ Inactivated SARS-CoV-2

Given that 5, 10, 15, and 20% Si_3_N_4_ were non-toxic to Vero cells, antiviral testing at these concentrations was performed. SARS-CoV-2 virions were exposed to Si_3_N_4_ at these concentrations for 1, 5, or 10 minutes. Following Si_3_N_4_ exposure, the infectious virus remaining in each solution was determined through plaque assay. Virus processed in parallel but only exposed to cell culture media contained 4.2 x 10^3^ PFU/mL. SARS-CoV-2 titers were reduced when exposed to all concentrations of Si_3_N_4_ tested (Figures 4A and B). The inhibition was dose-dependent with SARS-CoV-2 exposed for 1 minute and 5% Si_3_N_4_ having reduced viral titers by ~0.8 log_10_, 10% Si_3_N_4_ by ~1.2 log_10_, 15% Si_3_N_4_ by 1.4 log_10_, and 20% Si_3_N_4_ by 1.7 log_10_ (Figure 4A). Similar results were observed with the 5 and 10 minute samples. This reduction in viral titers corresponded to 85% viral inhibition at 5% Si_3_N_4_, 93% at 10% Si_3_N_4_, 96% at 15% Si_3_N_4_, and 98% viral inhibition at 20% Si_3_N_4_ (Figure 4B). Higher Si_3_N_4_ concentrations for longer times resulted in increased inhibition – leading to 99.6% viral inhibition at 20% Si_3_N_4_ and 10 minute exposure (Figure 4B). These data indicate that Si_3_N_4_ has a strong antiviral effect against SARS-CoV-2.

**Figure 4.**
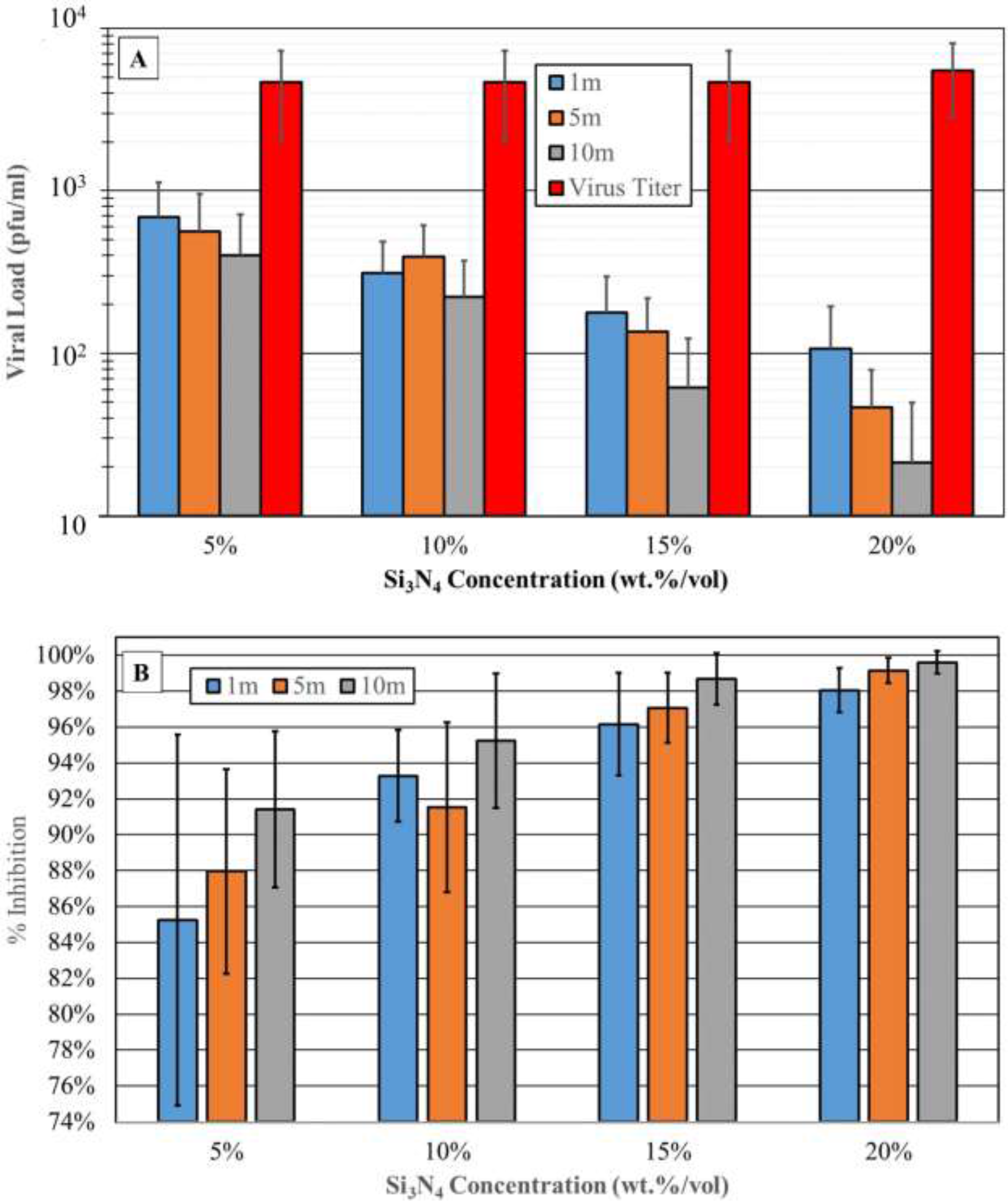
Silicon nitride antiviral activity testing. Silicon nitride at concentrations of 5, 10, 15, and 20 wt.%/vol were incubated with virus diluted in cell culture media for 1, 5, and 10m. At each timepoint, the samples were centrifuged, and the supernatant was collected and filtered through a 0.2um filter. The clarified supernatant was used to perform plaque assay in duplicate (neat through −5 dilutions were plated); (A) Data are expressed as PFU/mL. N=4 (two biological replicates and two technical replicates); (B) Data are expressed as % inhibition, with the virus only control (media only) set to 0%.

## 4. DISCUSSION

The remarkable finding in the present study is that a one-minute exposure to a 5% solution of Si_3_N_4_ resulted in 85% inactivation of SARS-CoV-2, while Vero cell viability was minimally impacted even after a 48 hour exposure to a 20% concentration of the same material. This finding is consistent with previous work showing that Si_3_N_4_ exerts a fortuitous dual effect, whereby the material can inactivate viruses^12, 26^ and inhibit bacterial biofilm formation^29^ without negatively impacting mammalian cell viability. The present study is also consistent with a recent investigation that showed the rapid inactivation of SARS-CoV-2 upon exposure to 15% (w/v) Si_3_N_4_,^12^ and related data showing that the antiviral effect of Si_3_N_4_ may be generally applicable to other single-strand RNA viruses such as Influenza A, Feline calicivirus, and Enterovirus.^26^ Taken together, these studies suggest that Si_3_N_4_ may be a suitable material platform for the development of fabrics for personal protective equipment such as masks, and also to manufacture commonly-touched surfaces where viral persistence may encourage the spread of disease.

The antiviral effect of Si_3_N_4_ is related to several probable mechanisms that have been hypothesized previously.^12^ One mechanism is the viral RNA fragmentation by reactive nitrogen species derived from a slow and controlled elution of ammonia from the surface of Si_3_N_4_; and the subsequent formation of ammonium, with the release of free electrons and negatively charged silanols in aqueous solution. Also, as reported previously, the similarity between the protonated amino groups, Si–NH_3_^+^ at the surface of Si_3_N_4_ and the N-terminal of lysine, C–NH_3_^+^ on the virus triggers a competitive binding that leads to SARS-CoV-2 inactivation.^12^ An advantage of Si_3_N_4_ is that these phenomena reflect a sustained mechanism of action secondary to a hydrolysis-mediated chemical equilibrium on the surface of the material, rather than a reliance on repeated applications required of commonly-used disinfecting agents.

While the antiviral effectiveness of Si_3_N_4_ is comparable to Cu, a historically-recognized viricidal agent, the use of Cu is limited by its cell toxicity.^30^ In contrast to Cu, ceramic implants made of Si_3_N_4_ that have the same composition as the material used in this study have shown successful clinical and radiographic outcomes even after three decades in the human body.^31^ An advantage of Si_3_N_4_ toward the potential development of antiviral and antibacterial surfaces is the versatility of the material; thus sintered Si_3_N_4_ has been incorporated into polymers, bioactive glasses, and even other ceramics to create composites and coatings that retain the favorable osteogenic and antibacterial properties of Si_3_N_4_.^32–36^

We recognize that this study has limitations; a powdered form of doped-Si_3_N_4_ was used, rather than the monolithic material employed as spine implants. While this study demonstrates that this powdered form of doped-Si_3_N_4_ is an effective antiviral agent, future studies will need to show that its viricidal efficacy is retained when compounded into or coated onto other materials such as polymers, paints, metals, fabrics, or ceramics. To be truly effective, the surface hydrolysis and release kinetics of *β*-Si_3_N_4_’s moieties will have to be optimized for each composite material and device. Additionally, further refinement of the current composition of doped *β*-Si_3_N_4_ may enhance its ability to release these moieties for the benefit of mammalian cells and the destruction of pathogens. Studies are also needed to identify whether simple physical contact with Si_3_N_4_ particles, or exposure to its soluble components, or both, are necessary for the antipathogenic effects observed in the present study.

## 5. CONCLUSIONS

In the present study, Si_3_N_4_ proved harmless to mammalian cells, while demonstrating a time- and dosedependent inactivation of SARS-CoV-2. While Si_3_N_4_ is not suitable for ingestion or inhalation, its antiviral activity, which is not limited to SARS-CoV-2, suggests that it may be a fortuitous platform to engineer surfaces and personal protective equipment to discourage viral persistence, and thereby control the spread of COVID-19 and other diseases.

## ACKNOWLEDGEMENT

The following reagent was deposited by the Centers for Disease Control and Prevention and obtained through BEI Resources, NIAID, NIH: SARS-Related Coronavirus 2, Isolate USA-WA1/2020, NR-52281.

